# Dimeric Transmembrane Structure of the SARS-CoV-2 E Protein

**DOI:** 10.1101/2023.05.07.539752

**Authors:** Rongfu Zhang, Huajun Qin, Ramesh Prasad, Riqiang Fu, Huan-Xiang Zhou, Timothy A. Cross

## Abstract

The SARS-CoV-2 E protein is a transmembrane (TM) protein with its N-terminus exposed on the external surface of the virus. Here, the TM structure of the E protein is characterized by oriented sample and magic angle spinning solid-state NMR in lipid bilayers and refined by molecular dynamics simulations. This protein has been found to be a pentamer, with a hydrophobic pore that appears to function as an ion channel. We identified only a symmetric helix-helix interface, leading to a dimeric structure that does not support channel activity. The two helices have a tilt angle of only 6°, resulting in an extended interface dominated by Leu and Val sidechains. While residues Val14-Thr35 are almost all buried in the hydrophobic region of the membrane, Asn15 lines a water-filled pocket that potentially serves as a drug-binding site. The E and other viral proteins may adopt different oligomeric states to help perform multiple functions.

## Introduction

The SARS-CoV-2 virus in the past few years has resulted in over 700 million COVID-19 infections and 6.9 million deaths worldwide.^1^ SARS-CoV-2 is an enveloped, positive-strand RNA virus that belongs to the coronaviridae family^2^ and is closely related to the SARS-CoV-1^3^ and MERS-CoV^4^ viruses that caused previous epidemics. Four proteins decorate the surface of SARS-CoV-2, including the membrane (M), envelope (E), nucleocapsid (N), and spike (S) proteins. The M, E, and S proteins are integral membrane proteins that are all reported to be homo-oligomers. The 75-residue E protein encodes a single transmembrane (TM) helix and has been proposed to form pentamers leading to a transmembrane pore that is capable of conducting ions.^5-11^ The SARS-CoV-2 virus is synthesized in the endoplasmic reticulum Golgi intermediate compartment (ERGIC) of mammalian cells.^8,11^ In the ERGIC and Golgi environment the N- and C-termini of the E protein are in the lumen and cytoplasm, respectively.^12^

Alam et al.^13^ have presented a functional pangenomic analysis revealing that SARS-CoV-1 and SARS-CoV-2 E proteins display two conserved key features, an N-terminal region with putative ion channel activity and a C-terminal PDZ-binding motif, that play key roles in inducing an inflammasome response leading to acute respiratory distress syndrome, a major cause of death from viral infections. Verdia-Bagguena et al.^14^ have suggested that ion conductance and cation selectivity are potentially controlled by the charge of the lipid membranes giving additional flexibility for viral reproductive conditions. While a portion of the E protein is in the viral membrane coat, most is located in intracellular transport sites of the host including the ER, Golgi, and ERGIC sites involved in viral assembly and budding.^15^ *In vitro* studies have shown that the removal of the E protein from recombinant coronavirus particles attenuates viral maturation and produces incompetent progenies.^16-18^ Consequently, the E protein is an important viral protein with multiple functions and possibly multiple conformational states in different environments.

Several solution NMR characterizations of the SARS-CoV-1 E protein in detergent micelles resulted in pentameric structures forming a TM hydrophobic pore that has been identified as an ion channel.^11,19,20^ An initial solid-state NMR (ssNMR) study of the TM domain (residues 8–38) in lipid bilayers of the SARS-CoV-2 E protein assumed this pentameric model, but added high-resolution structural detail for the TM helices (Protein Data Bank entry 7K3G).^21^ In 2021, a solution NMR study^22^ of the full-length SARS-CoV-2 E protein in detergent micelles did not challenge this oligomerization state and defined the secondary structure, revealing a putative transmembrane helix comprising residues 8-43 and a cytoplasmic helix comprising residues 53-60. In addition, a one-dimensional (1D) oriented-sample (OS) ssNMR spectrum in DMPC lipids was interpreted as indicating a large tilt angle, 45°, of the transmembrane helix. Although ion conductance through a pentameric assembly of E constructs that include the TM sequence has been demonstrated in various membrane mimetic environments using a large concentration gradient and/or membrane potential, whether the E protein has *in vivo* channel activity is still an open question.

For atomic-level structural studies of small membrane proteins in a native-like environment, ssNMR has clear advantages over other structural techniques such as X-ray crystallography, solution NMR, and cryo-electron microscopy (EM). Crystallization is only possible when contacts between protein molecules exist in a crystal lattice – for small TM proteins this often leads to structural distortions.^23^ Cryo-EM is growing as a technique for structural characterization of large protein assembles, but is not applicable to small proteins embedded in lipid bilayers. Solution NMR would also be a good approach if it was performed with the protein in nanodisc preparations.^24,25^ However, most solution NMR studies of membrane proteins have been performed in detergent micelles that can also lead to distorted structures.^23^ Taken together, with a sample in an extensive lipid bilayer environment, ssNMR is a technique that is capable of high-resolution structural characterization for small membrane proteins in a native-like environment.

We report here a SARS-CoV-2 E protein TM structure in planar lipid bilayers that results in a symmetric dimer that has not been previously characterized. Our extensive work on different E TM constructs with SDS-PAGE electrophoresis (Figure 1) and ssNMR contradicts the pentameric conclusion of prior characterizations. The dimeric structure of the SARS-CoV-2 E TM domain (E_12-37_) in a liquid-crystalline lipid bilayer environment clearly does not have the ability to conduct ions as previously observed for the pentameric characterizations. However, like the PDB 7K3G structure and multiple structural models, the tilt of the helices in our dimer in the lipid bilayer is small. In addition, the dimer is also homomeric, such that in our ssNMR spectra of singly labeled sites, only single resonances are observed. But, unlike the pentameric structures where each monomer has two different interaction surfaces with neighboring monomers, our structure has only a single interaction surface for each monomer, resulting in a symmetric dimer. The E protein is present in the viral coat at low copy numbers,^26^ where there is no evidence for channel formation. Potentially the dimeric structure characterized here represents the native structure of the E protein in the viral coat.

**Figure 1.**
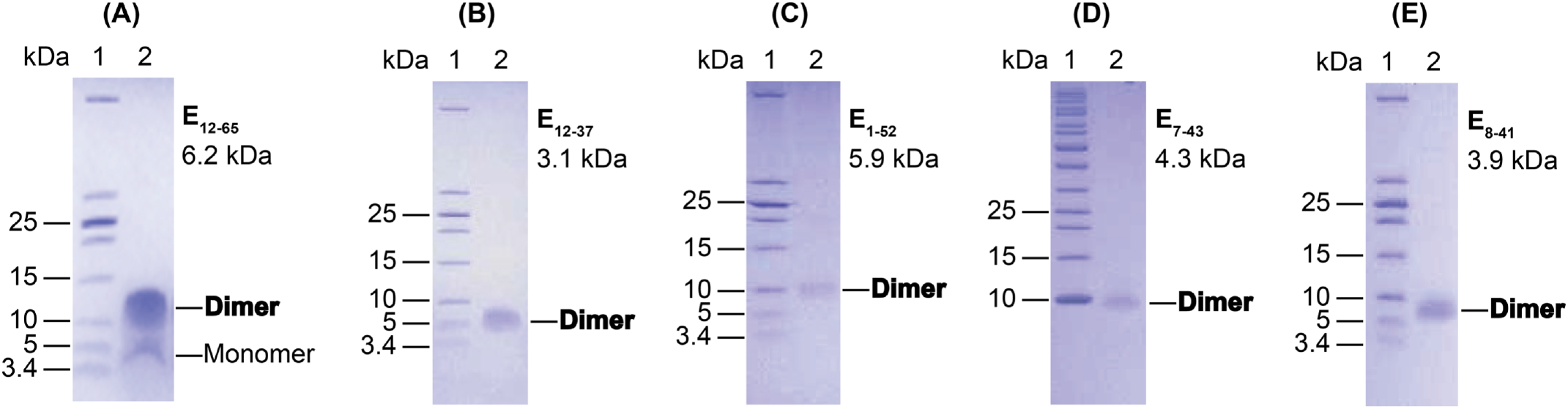
SDS-PAGE gels of purified E constructs. (A) E_12-65_. (B) E_12-37_. (C) E_1-52_. (D) E_7-43_. (E) E_8-41_. All of these constructs included leading Ser-Asn-Ala residues from the TEV cleavage site (Figure S1). The molecular weight of each construct is shown below its designation. The observed bands for monomer and dimer next to lane 2 in each gel are indicated. All display dominant bands for the dimer.

## Results

### TM sequence-containing E constructs form a stable dimer

We prepared a variety of E constructs (Figure S1), including E_1-52_, E_7-43_, E_8-41_, E_12-37_, and E_12-65_ that expressed well enough for SDS-PAGE electrophoresis (Figure 1). All of these constructs contained the TM sequence and ran primarily as dimers in our SDS-PAGE experiments, Importantly, there was no evidence of pentamers in the SDS-solubilized environment of these gels, the frequently suggested E oligomeric state in the literature. We used a ladder that included low molecular weight markers (down to 3.4 kD). Without the low molecular weight markers (Figure S2), the dimer band could be easily misidentified as a monomer.

We proceeded to prepare oriented samples of these E constructs in POPC/POPG bilayers (4:1 molar ratio), and achieved the highest alignment efficiency with E_12-37_. Our subsequent structural characterization thus focused on this E construct.

### E_12-37_ forms a highly regular TM helix with a small tilt angle

Figure 2A displays the 1D ^15^N ssNMR spectrum of a uniformly ^15^N labeled E_12-37_ OS with the lipid bilayer normal parallel to the magnetic field. The signal intensities spread over a very limited anisotropic ^15^N chemical shift range centered around 220 ppm, demonstrating a well-aligned sample. The central chemical shift is close to the σ_33_ element of the backbone ^15^N chemical shift tensor, indicating that the TM helical axis is close to the bilayer normal. The broadening on the low-field side of the spectra can be largely accounted for by the range of σ_33_ rigid-limit values (230-215 ppm) for the amino acid composition of E_12-37_. On the other hand, the broadening on the high-field side is more substantial, down to 175 ppm, suggesting that there is a small dispersion of amide ^15^N tensor orientations relative to the bilayer normal. A small tilt of the helical axis with respect to the bilayer normal is required to account for this dispersion. Another contributing factor is a small variation in the normal directions of the bilayers aligned between thin glass slides, typically no more than 2-3°.

**Figure 2.**
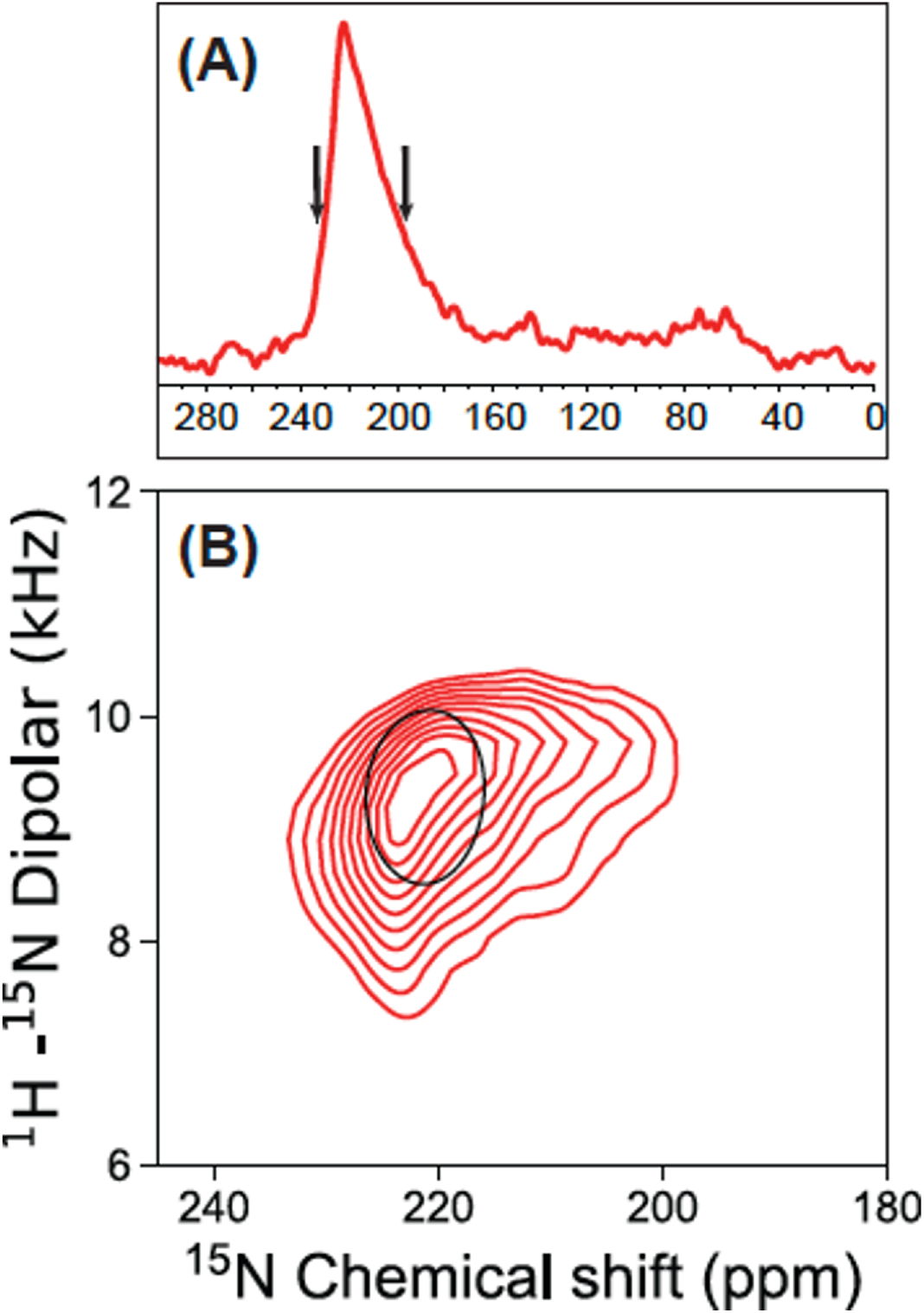
ssNMR spectra of uniformly ^15^N labeled E_12-37_ in aligned POPC/POPG bilayers. (A) 1D ^15^N spectrum. (B) 2D PISEMA spectrum of the same sample, with contours down to the spectral level as shown by the arrows on the 1D spectrum. Superimposed is a PISA wheel for a helix tilt angle of 6° with respect to the bilayer normal, calculated using a chemical shift tensor with σ_11_ = 54.5 ppm, σ_22_ = 82.8 ppm, σ_33_ = 227.1 ppm.^28^ The amino acid sequence for E_12-37_ with its N-terminal SNA tag is: SNAL_12_IVNSVLLF_20_LAFVVFLLVT_30_LAILTAL_37_.

Figure 2B displays a 2D spectrum of the same sample, obtained using the polarization inversion spin exchange at magic angle (PISEMA) pulse sequence^27^ that correlates the ^15^N anisotropic chemical shift with the ^1^H-^15^N dipolar coupling. Contours in the PISEMA spectrum are displayed only down to the level of the black arrows shown in the 1D chemical shift spectrum (Figure 2A), due to a more limited signal-to-noise ratio in the 2D PISEMA spectrum. The contours span the range of chemical shifts from 233-200 ppm. Importantly, this range encompasses all of the resonance frequencies for the amino acid specific labeled spectra presented below.

In the ^1^H-^15^N dipolar dimension, the PISEMA contours span a range from 7.5 to 10.4 kHz. These large values limited to a narrow range again show that the entire peptide forms a highly regular helix with a small tilt angle. There is not even a single residue at the peptide termini that has significantly reduced anisotropic chemical shift or dipolar coupling that could result from fraying.

The backbone ^15^N resonance frequencies in PISEMA spectra are inherently dependent on the orientations of the ^15^N chemical shift and ^1^H-^15^N dipolar tensors relative to the magnetic field, which is parallel to the bilayer normal. The orientations of the tensors in turn are determined by the orientation of the peptide plane. For water-soluble helices, the angle between the peptide plane and the helical axis is approximately 12°, meaning that even when the helical axis is aligned with the magnetic field, neither the ^1^H-^15^N dipolar vector nor the ^15^N σ_33_ element, which are both in the peptide plane, are parallel to the magnetic field. However, for TM helices the peptide plane tilt angle is reduced to 6° with respect to the helical axis, due to the repulsive interaction between the carbonyl oxygens and fatty acyl environment of the lipid bilayer compared to an attractive aqueous environment for the carbonyl oxygens.^28^ ^29^ Consequently, for a TM helix aligned with the magnetic field, the PISEMA spectrum would reflect the backbone N-H bond vectors at a fixed angle of ∼6° with respect to the magnetic field. When the helix is tilted with respect to the magnetic field, the range of angles increases. For example, for a helix tilted at 6°, the angles of the N-H bond vectors with respect to the magnetic field would range from ∼0° to ∼16°, depending on the self-rotation of the helix. To determine the tilt and rotation of the TM helix, numerous amino acid specific ^15^N labeled samples of E_12-37_ were prepared for PISEMA spectroscopy.

Figure 3 displays six sets of 2D PISEMA spectra for six E_12-37_ preparations using amino acid specific ^15^N labeling. With these samples, 24 of the 26 residues of E_12-37_ are labeled. The residues that were not labeled were Asn15 and Ser16. All of the observed resonances fall in a very small region that overlaps with the PISEMA spectrum of the uniformly ^15^N labeled sample. Together, the PISEMA spectra indicate that the entire E_12-37_ sequence, including Asn15 and Ser16 and even Ala11 (part of inserted TEV cleavage site and not an E residue), forms an α-helix. As illustrated in Figure S3 for the Val ^15^N labeled sample, there is no other spectral intensity other than those shown in the spectra of Figures 2 and 3. Indeed, the observed resonances for the entire sequence fall in a very narrow region and overlap with a polarity index slant angle (PISA) wheel drawn for a helix having a 6° tilt with respect to the bilayer normal. A PISA wheel presents the predicted PISEMA resonances as the peptide plane traces an ideal α-helix trajectory that has a preset tilt angle.^30,31^

**Figure 3.**
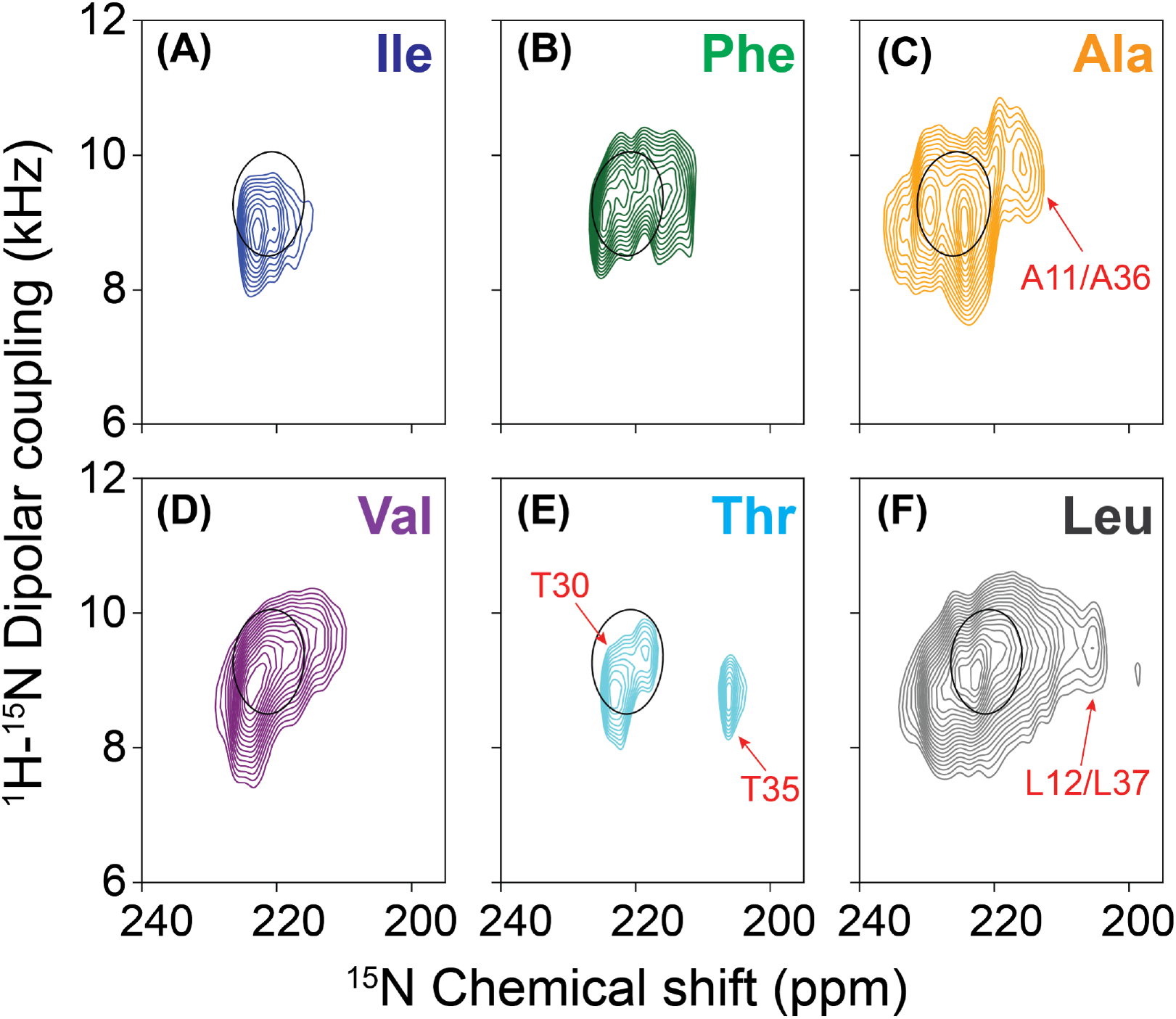
PISEMA spectra of ^15^N labeled E_12-37_ peptides in POPC/POPG bilayers. (A) ^15^N-labeling at Ile13 and Ile33. (B) ^15^N-labeling at Phe20, Phe23, and Phe26. (C) ^15^N-labeling at Ala11, Ala22, Ala32, and Ala36. D) ^15^N-labeling at Thr30 and Thr35. (E) ^15^N-labeling at Val14, Val17, Val24, Val25, and Val29. (F) ^15^N-labeling at Leu12, Leu18, Leu19, Leu21, Leu27, Leu28, Leu31, Leu34, and Leu37. Superimposed on the spectra is a PISA wheel calculated at a tilt of 6°. For the Ala spectrum, isotopic labeling included the Ala residue (denoted as Ala11) from the TEV cleavage site, and the calculated PISA wheel used a somewhat larger σ_33_ of 232.1 ppm.

The chemical shift linewidths observed in our oriented samples were typically less than 5.0 ppm over the total chemical shift range of 170 ppm. The narrow linewidths document excellent alignment of our samples resulting in high-resolution structural restraints. Normally, we would attempt to assign resonances to a PISA wheel pattern having 3.6 resonances per turn of a helical wheel, but the tilt of the helix here is so small that even with the excellent linewidths of the individual resonances of a given amino acid type it is not possible, with a few exceptions noted below, to make such assignments. For amino acid type labeled preparations such as Val with more than two residues labeled, the dispersion is too small to generate spectral resolution of the individual resonances. However, the resonances in the first and last turns of the helix display some additional dynamics due to each peptide plane having a single backbone hydrogen bond. The increased dynamics shifts the anisotropic chemical shift resonances slightly upfield and also reduces spectral intensity, such as that for Thr35. As a result, the Thr30 and Thr35 residues can be assigned to the resonances at 219 and 205 ppm, respectively. Likewise, the extra intensities in the upfield region of the Ala ^15^N labeled and Leu ^15^N labeled spectra can be tentatively assigned to Ala11, Leu12, Ala36, and Leu37, in the first and last turns of the helix. For the Ala residues, the tensor has a σ_33_ that is considerably greater than the other hydrophobic amino acids and consequently the entire spectral envelope is shifted downfield compared to the other residues by ∼5 ppm.^28^ The PISEMA spectra, especially the one from the ^15^N-Thr labeled sample, all point to the E_12-37_ structure being homomeric; i.e., corresponding residues (e.g., Thr30) in different subunits experience the same chemical environment.

The OS alignment efficiency of longer construct E_8-41_ was not as good as that of E_12-37_. In contrast to the PISEMA spectrum of Val ^15^N labeled E_12-37_, where all the resonances are around the PISA wheel for a 6° helix tilt, the corresponding spectrum of E_8-41_ has extra intensities below 150 ppm (Figure S3). These upfield intensities arise from unoriented materials as well as the dynamic residues 8-11 and 38-41 of aligned E_8-41_ that are outside the lipid bilayer, and constitute the so-called powder pattern.

### E_12-37_ forms a symmetric helix-helix dimer interface

Having defined the helix tilt by the OS experiments, we proceeded to define the interhelical interface by magic angle spinning (MAS) experiments. To obtain interhelical distance restraints, different amino acid type ^13^C-labeled E_12-37_ peptides were prepared and mixed in a 1:1 molar ratio. We utilized the ^13^C-^13^C correlations between two types of ^13^C-labeled amino acids to identify interhelical cross peaks, as the correlations through the dipolar recoupling at different mixing times provide distance measurements for up to 8 Å between two ^13^C nuclei.^32,33^

The labeling schemes were carefully chosen based on the primary sequence and helical wheel projection. Because the E TM sequence consists of mostly long sidechain residues, there were no obvious interaction motifs (such as Gly at *i* and *i* + 4 or *i* + 7 positions) to guide the labeling. We first chose Val and Leu residues as labeling sites, because all 5 Val residues locate within an arc of the helical wheel that spans 140°, while the 9 Leu residues populate throughout the whole helical wheel (Figure S4A). This choice could give a reasonable chance of obtaining interhelical cross peaks while still narrowing down the interhelical interface.

As shown in Figure 4A, for the ^13^C-Leu and ^13^C-Val mixture at a 600 ms mixing time, prominent interhelical cross peaks were observed for: LCα-VCα, LCβ-VCα, LCγ-VCα, LCα-VCβ, LCβ-VCβ, and LCγ-VCβ, which included all possible pairs of Val and Leu aliphatic carbons except for those at the branched termini, which are highly dynamic due to bond rotation. In particular, the Cα-Cα cross peak indicates a remarkably close approach between the helix backbones at the interface. When the mixing time was reduced to 300 ms (Figure 4B), we were still able to observe the LCα-VCα cross peak, suggesting a distance of ≾6.5 Å.^33^ With a further reduction of the mixing time to 100 ms, the interhelical cross peaks were no longer observable due to insufficient time for interhelical polarization transfer (Figure S5).

**Figure 4.**
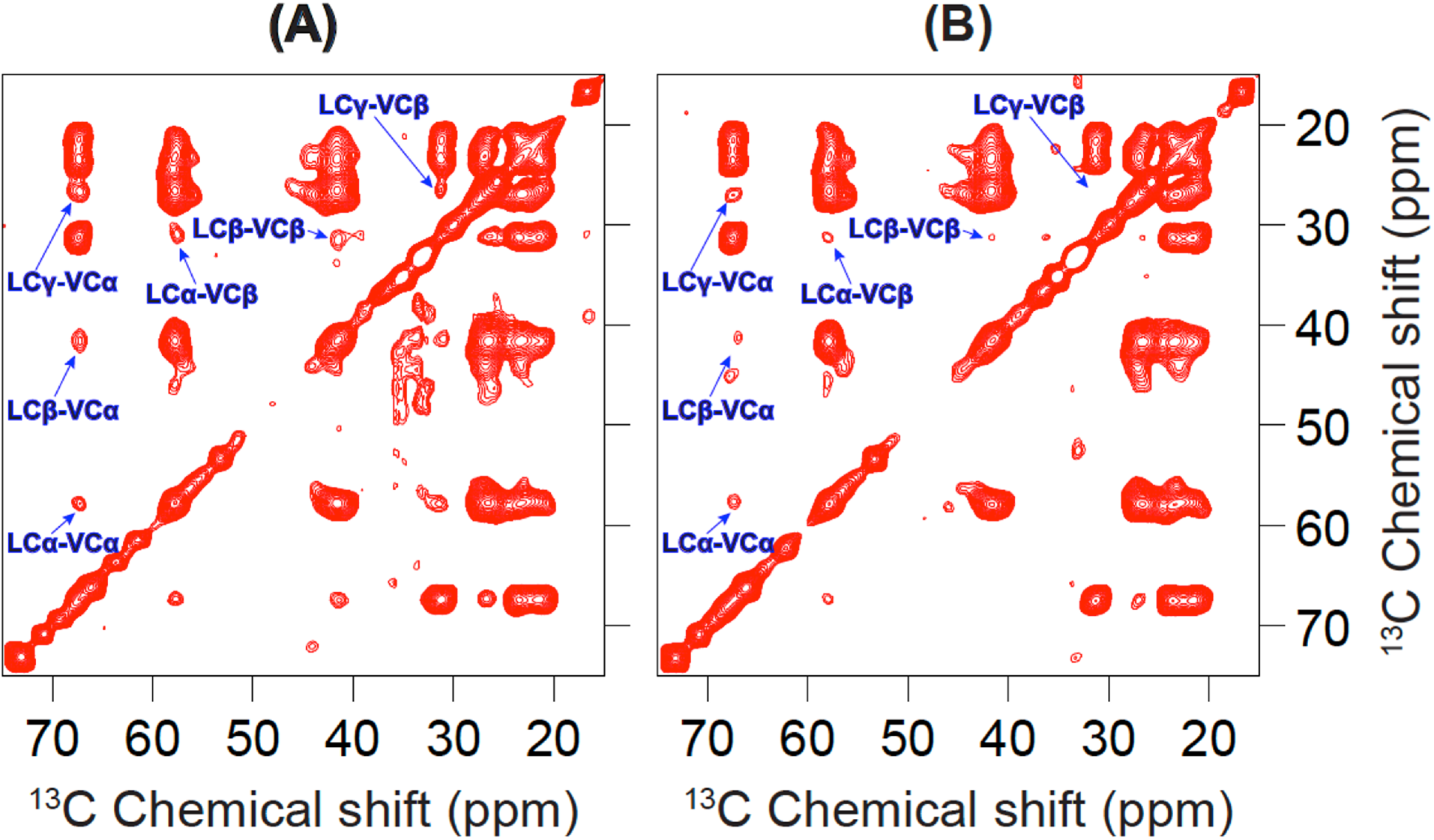
^13^C-^13^C correlation spectra of a 50:50 mixture of ^13^C-Leu labeled E_12-37_ and ^13^C-Val labeled E_12-37_ in POPC/POPG liposomes. Data obtained at mixing times of (A) 600 ms and (B) 300 ms. Interhelical cross peaks are assigned.

Because there are 9 Leu and 5 Val residues in the TM sequence, at first sight it seemed not possible to unambiguously assign the cross peaks to specific Leu-Val pairs. However, the Leu-Val pairs that are most likely to be at the interhelical interface are those with one residue at position *i* and the other at *i* ± 1. Indeed, the interface of a homo-oligomer must have *i* to *i* ± 1 contacts. In the E TM sequence, there are two *i* to *i* ± 1 Leu-Val pairs: Val17-Leu18 and Leu28-Val29. Given the strong intensities of the Leu-Val cross peaks and the fact that at most one quarter of the mixture could contribute to any cross peak, most likely both of these Leu-Val pairs generated cross peaks.

To further narrow down the possibilities of the interhelical interface, we introduced a V25M mutation and prepared a sample mixing the ^13^C-Val labeled mutant with the ^13^C-Met labeled mutant (Figure S6). No interhelical cross peaks were observed on the ^13^C-^13^C correlation spectrum of this sample at a mixing time of 600 ms, suggesting that Val24 and Val25 cannot be both present at the interhelical interface.

In our third MAS sample, the ^13^C-Leu labeled peptide was mixed with the ^13^C-Phe labeled peptide, because in the helical wheel the 3 Phe residues are located on a 120° arc opposite to the 140° arc spanned by the 5 Val residues (Figure S4A). As shown in Figure 5, no cross peaks were observed between the aliphatic carbons of Leu and Phe, and only a weak cross peak was observed between Leu Cγ and a Phe aromatic carbon. So the Cα atoms of all Leu-Phe pairs are far apart, but their long sidechains may reach each other within ∼8 Å.

**Figure 5.**
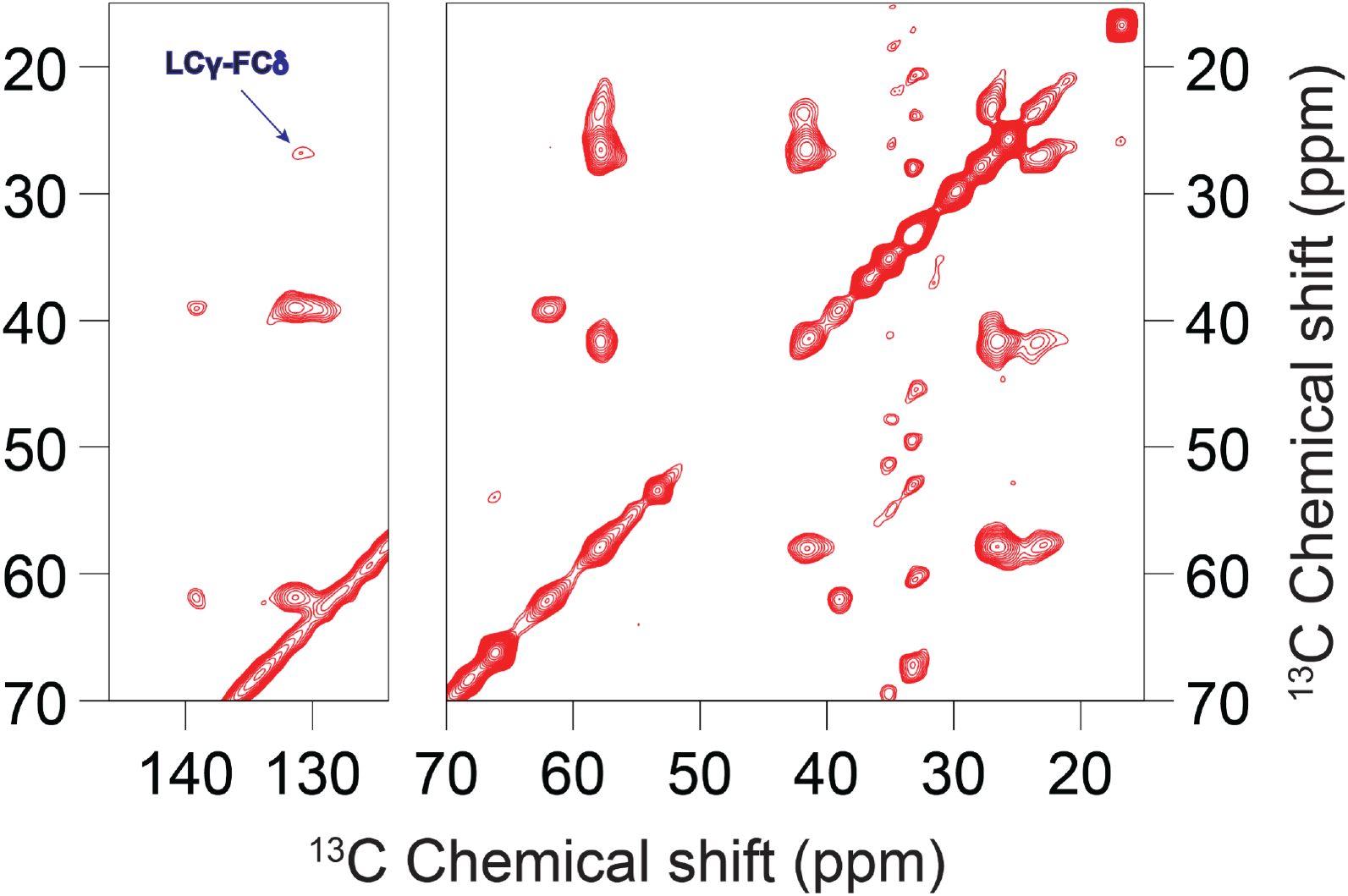
^13^C-^13^C correlation spectrum of an equimolar mixture of ^13^C-Leu labeled E_12-37_ and ^13^C-Phe labeled E_12-37_ in POPC/POPG liposomes at a mixing time of 600 ms.

We used the above restraints, both positive and negative, from the ^13^C-^13^C correlation spectra to uniquely define the interhelical interface, by scanning for all possible helix-helix arrangements. The observation that one side of the helical wheel (the “Val face”) is in the interface while the opposite side (the “Phe face”) is not in the interface strongly suggests a symmetric dimer interface. Scanning over helix rotation produced only a single dimer interface, shown in Figure S4B, that satisfies all the restraints. In this interface, Leu21 and Val25 are at the core, and Val17, Leu18, Leu28, and Val29 are at the periphery. In contrast, the interhelical interface of a pentamer structure is asymmetric and pairs the Val face with the Phe face (or portions thereof). Thus unavoidably at least one of the Phe Cα atoms is placed at the interface. For example, the solution NMR structure 5×29^19^ has the Phe23 Cα at the center of the interface. Our scanning of helix rotation produced a pentamer with an interface that minimizes the burial of Phe Cα atoms (Figure S4C), but even in this model, Cα-Cα cross peaks for both Val24-Val25 and Phe26-Leu28 pairs would be observable. This model is very similar to the ssNMR structure 7K3G, which, in addition to the Cα-Cα contacts of Val24-Val25 (6.4 Å) and Phe26-Leu28 (7.3 Å) that would give rise to observable cross peaks, has a close Phe26 Cδ – Leu27 Cγ distance (5.1 Å) that would generate a strong cross peak. These expectations are inconsistent with our ^13^C-^13^C correlation spectra (Figure S6 and Figure 5).

### Dimer structure of E_12-37_ refined in POPC/POPG membranes exposes Asn15 to a water-filled pocket for potential drug binding

Based on the well-defined helix tilt by the OS data and the well-defined dimer interface by the MAS data, we built an initial model by Xplor-NIH^34^ and refined the structure by restrained molecular dynamics simulations in POPC/POPG membranes (Figure 6). The resulting structure satisfies all the NMR restraints, including the helix tilt angle of 6°; Cα-Cα distances ≤ 6.5 Å for all Val17-Leu18 and Leu28-Val29 pairs; no Leu-Phe Cα-Cα distance < 9 Å; and an 8.2 Å distance between Phe26 Cδ and Leu28 Cγ, which can account for a weak cross peak between this pair of atoms (Figure 5). We repeated the refinement simulations 14 times; the resulting models are very close to each other, with backbone root-mean-square deviations of 0.7 ± 0.2 Å from model 1 (Figure S7A).

**Figure 6.**
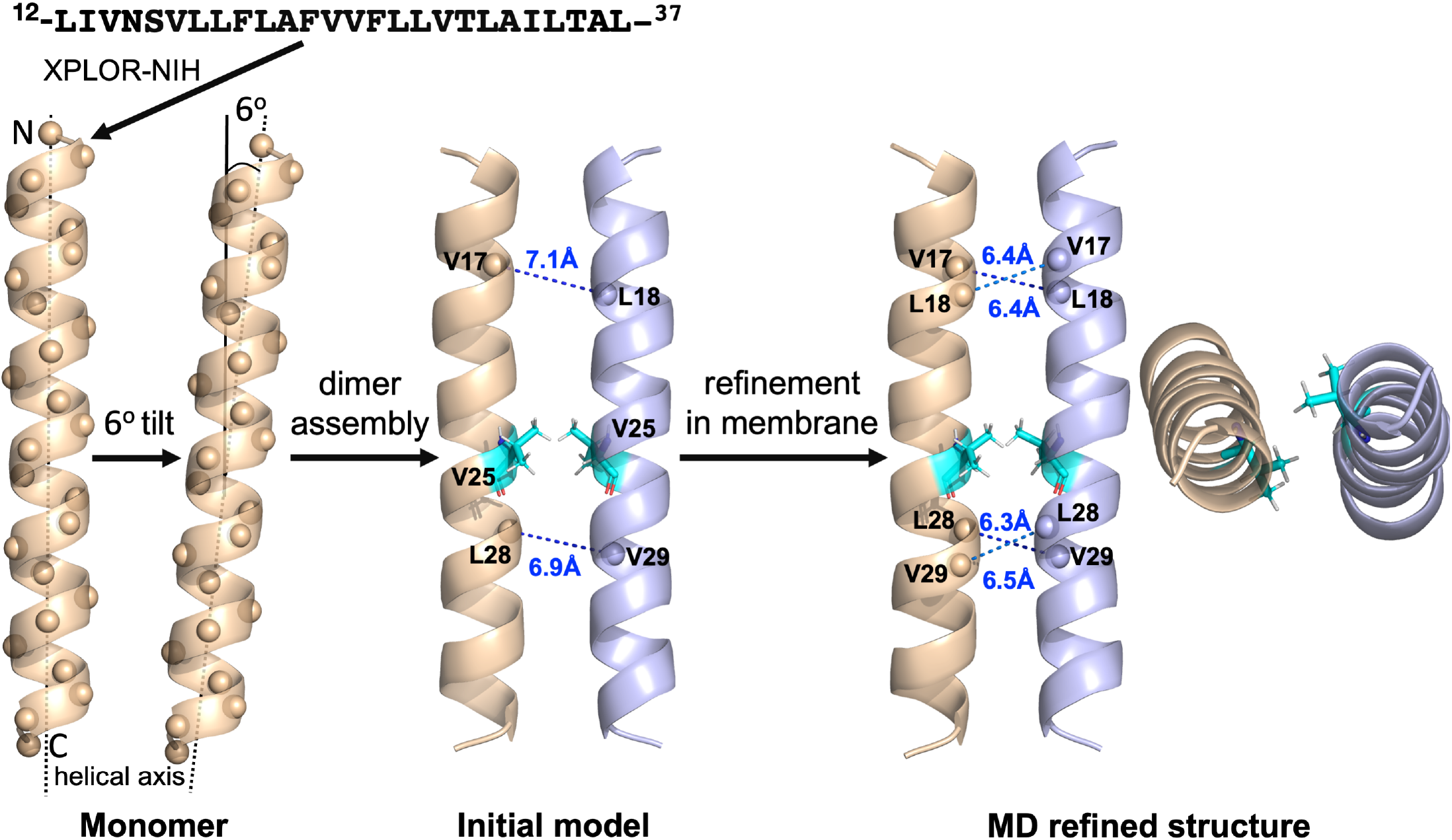
Refinement of the E_12-37_ dimer by restrained molecular dynamics simulations in POPC/POPG membranes.

The Cα atoms of residues Val14-Thr35 are positioned in the hydrophobic region of the membrane. The hydrophobic region, well-suited for the nonpolar sidechains among these residues, may pose a problem for the polar sidechains, of which there are four: Asn15, Ser16, Thr30, and Thr35. The sidechain hydroxyls of both Thr30 and Thr35 form hydrogen bonds with the backbone carbonyls four residues upstream (Figure S7A). On the other hand, Asn15 and Ser16 are close enough to the membrane surface such that water molecules push away the lipids to form hydrogen bonds with these polar sidechains (Figure S8A). The numbers of water molecules are only 3 to 4 around each Ser16 sidechain but approximately 15 around each Asn15 sidechain, generating a water-filled pocket that may serve as a drug-binding site (Figure S8B).

## Discussion

We have combined OS and MAS ssNMR, molecular modeling, and restrained molecular dynamics simulations to determine the dimeric structure of the SARS-CoV-2 E protein in liquid-crystalline lipid bilayers. Resonances in the PISEMA spectra are limited to a small region around 220 ppm in ^15^N anisotropic chemical shift and 9 kHz in ^1^H-^15^N dipolar coupling. Such narrowly limited large values in both dimensions clearly indicate that the entire sequence of E_12-37_ forms a highly regular helix with a small tilt angle. MAS ^13^C-^13^C dipolar restraints unequivocally define a symmetric interhelical interface, pointing to a dimer structure. The refined dimer structure of E_12-37_ has a helix tilt angle of only 6°, with an extended interface dominated by Leu and Val sidechains. Residues Val14-Thr35 are positioned in the hydrophobic region of the membrane, but a pocket in the membrane surface allows the Asn15 sidechain to access water. Asn15 has been shown to be the major site for binding the amiloride class of drugs.^22^ Because these molecules have a high tendency for lipid partition,^35^ they may bind to Asn15 from the lipid bilayer.

The first clear indication of dimers was provided by SDS-PAGE gels, where the E protein was solubilized by the detergent SDS. Although we did not directly determine the oligomeric state of E_12-37_ in lipid bilayers, dipolar restraints from ^13^C-^13^C correlation spectra allowed us to unequivocally define a symmetric, dimer interface. All previous solution NMR characterizations of the E protein in detergent micelles have produced pentameric structures;^11,19,20^ a recent ssNMR study also assumed a pentameric model.^21^ Our ^13^C-^13^C dipolar restraints specifically led to the elimination of pentameric models. Contrary to the small tilt angles in our dimer structure and in the pentamer structure of Mandal et al.,^21^ Park et al.^22^ suggested a tilt angle of 45° based on a 1D OS spectrum in DMPC lipids, ostensibly to accommodate a long helix (extending to Glu8 on the N-terminal side) that was determined by solution NMR in detergent micelles. In our ssNMR structure, residues Leu11 and Ile12 are positioned in the membrane interfacial region; though the helix could extend further into the N-terminal side, these residues (M_1_YSFVSEETGT_11_) would be in the extraviral aqueous environment.

The function of the SARS-CoV-2 E protein has often been described as an ion channel in a pentameric state. This putative activity is difficult to understand because an 18-residue long stretch, from Val17 to Leu34, of the TM sequence contains just one weakly hydrophilic residue, Thr30. The resulting highly hydrophobic pore would not be favorable for ion conductance or even for water exposure. It is unclear why such a hydrophobic pore would have evolved for ion conductance. Although ions may be induced to flow through such an unfavorable pore with high concentration gradients, this conductance activity is not likely to be relevant in the viral envelope where relatively few copies of the protein are found.^26^ Indeed, the literature has not suggested such activity in the viral envelope.

The TM domains of E orthologs in related viruses have been found to be required for both virus assembly and release.^36,37^ Alanine insertions in the TM helix of the mouse hepatitis coronavirus (MHV) A59 E protein significantly crippled virus production, and the effect was attributed to the disruption of the alignment of polar sidechains on the surface of the TM helix.^36^ Polar residues Gln15, Cys23, and Thr27 in this E protein are all positioned on one face of the TM helix, and the authors suggested that they may participate in interactions with the TM domain of the M protein. Interestingly, the polar face of the MHV E protein aligns with what we have referred to as the Phe face in the SARS-CoV-2 E protein (Figure S4B). This face is away from the dimeric interface and thus available for interacting with other proteins; it also harbors three polar residues of the SARS-CoV-2 E protein: Asn15, Ser16, and Thr30. If this face indeed participates in interactions with the M protein, then the inhibitory effects of the amiloride class of drugs may have an alternative mechanism. Instead of acting as a channel blocker, they inhibit E-M protein binding.

Unlike many TM helical domains that are stabilized by close helix-helix packing afforded by short sidechains (Gly or Ala) at *i* and *i* + 4 or *i* + 7 positions, the E dimer structure determined here is stabilized by an extended interface comprising mostly of Leu and Val sidechains. This interface has some resemblance to those of leucine zippers, which are distinguished by a Leu residue at every seventh position. Each Leu sidechain at position *i* of one helix interacts with the corresponding Leu sidechain (at *i*’) and two other hydrophobic sidechains at *i*’ – 3 and *i*’ + 4 on the opposite helix. Salt bridges between oppositely charged sidechains at *i* – 4 and *i*’ + 1 or at *i* + 1 and *i*’ – 4 provide further stabilization. These charged sidechains also contribute to solubility in water. The E protein has Leu residues at positions 21 and 28; Leu21 on one helix of the dimer indeed interacts with Leu18, Leu21, and Val25 on the opposite helix (Figure S4B), but that is where the resemblance to a leucine zipper ends. Supercoiling of leucine zippers reduces the helical periodicity to 3.5 residues per turn, whereas straight helices in TM domains have a periodicity of 3.6 residues per turn. As a result, Leu28 does not reach the interface as deeply as Leu21 and forms hydrophobic interactions with Val25 and Val29 on the opposite helix. If Leu28 reached the same position as Leu21 in the interface, it would interact with Val25, Leu28, and Aal32 on the opposite helix.

The dimer structure of E_12-37_ was determined from samples with a 1:25 protein-to-lipid molar ratio. With increasing protein-to-lipid ratio, dimers might form clusters (as found for the TM domain of the influenza M2 proton channel^38^) or perhaps even reorganize into higher oligomers. In nanodiscs, small viral proteins have been observed to populate a range of oligomeric states.^39^ It may well be that in the ERGIC where the virus is assembled and the E protein is present at high concentrations,^15^ it forms pentamers, but in the mature virus where it is present at a low copy number,^26^ it only forms dimers. Viral proteins often perform multiple functions. The change in oligomeric state may facilitate that.

## Materials and Methods

### Gene cloning

The gene sequences of all E fragments were based on the SARS-CoV-2 isolate Wuhan-Hu-1 (NC_045512). As illustrated in Figure S1A, four fragments E_12-37_, E_1-52_, E_7-43_, and E_8-41_ were separately cloned into the pTBMalE vector ^40^ following our previous work on influenza A M2 protein.^41 42^ For the fifth fragment E_12-65_, the pTBGST vector ^40^ was used as illustrated in Figure S1B. All gene constructs contained a 6 × His-tag for protein purification and a TEV cleavage site (ENLYFQSNA) for fusion protein removal. Cysteines in the E fragments were mutated to serine for E_1-52_, E_7-43_, and E_8-41_ and to alanine for E_12-65_. Cloned gene constructs were all verified by DNA sequencing.

### Protein expression and purification

*E. coli* BL21(DE3)RP cells were used to express all E constructs. Similar expression conditions were used for vectors containing GST and MBP fusion proteins. In short, 100 ml each of overnight Luria Broth (LB) culture was evenly divided into two 1-L LB media and grew at 37 °C with vigorous shaking to an OD_600_ of ∼0.8. The cells were spun down and washed with 30 ml of sterile M9 media. They were then resuspended in 500 ml of M9 media supplemented with 100 mg of a desired ^15^N-labeled or ^13^C-labeled L-amino acid (Cambridge Isotope Labs, Inc. or Sigma-Aldrich, Inc.) as well as the remaining 19 unlabeled amino acids. Cultures grew at 37 °C for another 20 min and over-expression was induced for 1.5 hr with 0.4 mM IPTG. Upon over-expression, the cells were harvested and resuspended in 60 ml of buffer (20 mM Tris-Cl, pH 8.0, 500 mM NaCl) prior to storage at -80 °C. For the expression of uniformly ^15^N-labeled proteins, the sole nitrogen source in the M9 minimal media was replaced with ^15^NH_4_Cl, and the induction time was increased to 8 hr.

Cells were thawed in a 37 °C water bath prior to adding 0.25 mg/ml of lysozyme and 5 units Benzonase nuclease (Millipore Sigma) per ml of cells. Cells were fully lysed by sonication (Fisher-100) using three cycles of 1.5 min pulsing and 1.5 min rest. 3% Empigen (Millipore Sigma) was then added and the tube was rotated at 4 °C for 2-3 hr prior to centrifugation at 10,000 g for 30 min. The supernatant was mixed with 20 ml nickel affinity resin (Qiagen) and rotated at 4 °C overnight.

The column was washed initially with buffer (20 mM Tris-Cl, pH 8.0, 500 mM NaCl, 3% Empigen, and 3 mM imidazole) and then for the second and third wash with buffer containing reduced Empigen (1.5% and 0.75% of Empigen, respectively). For MBP fusion protein purification, the final wash used a buffer without Empigen (20 mM Tris-Cl pH 8.0, 500 mM NaCl). While for GST, the Empigen in the final wash was replaced with 0.2% DPC (20 mM Tris-Cl pH 8.0, 500 mM NaCl, 0.2% DPC), until the UV absorbance returned to baseline. The protein was eluted with buffer (20 mM Tris-Cl pH 8.0, 500 mM NaCl, and 200 mM imidazole). Generally, 160-200 mg of MBP fusion protein and ∼80 mg of GST fusion protein were obtained per L of culture, respectively.

The MBP fusion protein was digested with Tobacco Etch Virus (TEV) protease at a protein-to-protease molar ratio of 1:2. The mixture was precipitated by adding 6% trifluoroacetic acid. The protein was then pelleted and washed with ddH_2_O twice, 100 mM Tris-Cl pH 8.0 twice, and finally ddH_2_O. The wet pellet was placed in a vacuum chamber overnight to remove any water content. The following day, methanol was added to the dried protein powder to extract the R fragment. Upon centrifugation at 15,000 g, methanol containing the E fragment was carefully removed and placed under a stream of nitrogen gas to remove the bulk methanol. Residual methanol was removed by vacuum overnight and the E fragment was ready for NMR sample preparation. The typical yield of E_12-37_ was ∼ 8–10 mg/L of culture.

### SDS-PAGE electrophoresis

The oligomeric states of the E fragments were assessed using SDS-PAGE electrophoresis. SDS buffer (2% SDS, 100 mM DTT, 10% glycerol, 100 mM Tris-HCl, and 0.01% bromophenol blue dye) was added to each E fragment preparation. After brief vortexing, the sample was loaded onto a 12% SDS-PAGE gel (Tris-Tricine). The sample was electrophoresed at room temperature at a constant voltage of 200 V for 40 min. After completion, the SDS-PAGE gel was stained with EZ-blue (Bio-Rad).

### NMR sample preparation

For oriented samples, lipid films were made using 25 mg of chloroform-dissolved POPC/POPG lipids (Avanti Polar Lipids, Inc.) in a 4:1 molar ratio. Most of the chloroform was removed by passing nitrogen gas over the sample. The remaining chloroform was removed in a vacuum chamber overnight. 1 ml of trifluoroethanol (TFE) was added to 4 mg of an E fragment and 2 ml of TFE was added to 25 mg of lipid film separately. They were then mixed at a 1:25 protein-to-lipid molar ratio and the solution was adjusted to pH 7.5 by adding a few μl of 0.5 N NaOH. The mixture was spun at 90 rpm for 2-3 hours and then spread evenly on 25 thin glass slides (5.7 mm × 10 mm × 0.04-0.06 mm, Matsunami Glass, Inc.). The glass slides were left in the vacuum chamber overnight to remove any trace of TFE, and then transferred to a hydration chamber (23 °C at 98% relative humidity via a saturated solution of potassium sulfate) for 2 hours. All glass slides were stacked together into a glass cell (6.3 mm × 4.3 mm × 18 mm, New Era Scientific Inc.) and sealed with a homemade plastic plug and beeswax (Hampton Research Inc.). MAS samples were prepared in a similar manner. Instead of spreading over glass slides, the hydrated proteoliposomes were packed into 3.2 mm thin-walled MAS rotors (Revolution NMR LLC).

### Solid-state NMR spectroscopy

All OS ssNMR experiments were performed on a Bruker NEO 600 NMR spectrometer with a Larmor frequency of 600.13 MHz for ^1^H. A custom-built low-electrical-field static probe with a rectangular coil was used. 1D cross-polarization and 2D PISEMA spectra were collected at 22 °C. The initial ^1^H 90° pulse strength was set to 50 kHz. The cross-polarization contact time was set to 1000 ms. A 50 kHz B_1_ field was used for both Lee–Goldburg irradiation and the Spinal-64 decoupling scheme during acquisition.^43^ The frequency jumps used to satisfy the Lee–Goldburg off-resonance condition were set to ±40.8 kHz. The number of t1 increments was 16; the number of scans per t1 increment was 3072 to 5128 depending on sample sensitivity.

For MAS NMR, a custom-built 3.2 mm low-E double-resonance MAS probe^44^ was used on a Bruker Avance 600 NMR spectrometer. The sample spinning rate was maintained at 10 kHz ± 3 Hz and the reading temperature was set to 228 K. The ^13^C magnetization was enhanced by cross polarization with a contact time of 1 ms. The ^1^H 90° pulse length was 3.2 μs before cross polarization. A phase alternated recoupling irradiation scheme,^45^ replacing the traditional dipolar assisted rotational resonance pulse sequence,^46,47^ was utilized with different mixing times (up to 600 ms) to identify interhelical cross peaks. The ^13^C chemical shifts were referenced to the carbonyl carbon resonance of glycine at 178.4 ppm relative to tetramethylsilane.

### Search for oligomer models

An ideal α-helix was built for the 26 Cα atoms of E_12-37_ using the following parameters: 3.6 residues per turn (corresponding to a 100° increment per residue on the helical wheel), 2.3 Å radius, and 1.5 Å rise per residue. The sequence of E_12-37_ was assigned to the Cα atoms on the helix, with an arbitrary initial rotation of the helix. A duplicate was then placed to the right of the first helix, with a 5.1 Å spacing between the rims of the helices.^48^ The second helix was also self-rotated by an angle appropriate for a given oligomeric state: 180° for a dimer and 72° for a pentamer.

The rotation angles of the two helices were scanned from 0 to 360° with a 10° interval. For each helix rotation angle, interhelical Cα-Cα distances were calculated to identify possible DARR contacts, with a cutoff of 7.2 Å.^33^ Models were filtered according to the following criteria: (1) no DARR contacts between residues 24 and 25; (2) no DARR contacts between any Leu-Phe pairs; (3) at least one DARR contact between a Leu-Val pair.

### Structure refinement for E_12-37_ dimer

A monomer model of E_12-37_ was built as an α-helix using Xplor-NIH,^34^ with backbone phi and psi angles assigned in the ranges -65° to -60° and -45° to -40°, respectively. The monomer was oriented with a 6° tilt angle with respect to the *z* axis; a duplicate was rotated to generate a dimer with C2 rotational symmetry. The dimer was subject to optimization by simulated annealing, with the following harmonic restraints: (1) backbone phi and psi angles, restrained to the above assigned values; (2) backbone *i* to *i* + 4 hydrogen bonding, with O-N distance restrained to 3.0 Å and O-H distance restrained to 2.0 Å; (3) interhelical Cα-Cα distances between Val17-Leu18, Leu28-Val29, and Val25-Val25 pairs, restrained to 6.0 Å. During the simulated annealing, the temperature was cooled from 3500 K to 25 K in 1000 steps, and the force constants for the restraints were ramped up from 5 to 1000 kcal/mol rad^-2^ for phi and psi angles and from 2 to 30 kcal/mol Å^-2^ for distances. The annealing simulation was repeated 100 times; a dimer model was selected based on considerations of the helix tilt angle (∼ 6°) and the distances between Val17-Leu18 and Leu28-Val29 pairs (∼ 7 Å).

Molecular dynamics simulations for the refinement of the initial model were run in NAMD^49^ with the CHARMM36 force field.^50^ Using the CHARMM-GUI web server,^51^ the initial model was placed in a POPC/POPG bilayer (4:1 ratio; 100 lipids per leaflet). The system was solvated with 8453 TIP3P water molecules, along with Na^+^ and Cl^-^ ions for charge neutralization and for generating a salt concentration of 30 mM, resulting in a simulation box of 83.6 × 83.6 × 81.5 Å^3^. After energy minimization (10000 cycles of conjugate gradient), simulations were run in four segments, with the first two at constant temperature and volume and the second two at constant temperature and pressure, for 125, 125, 125, and 250 ps, respectively. During these simulations, restraints on the lipids were gradually reduced, but positional restraints on the protein backbone heavy atoms were maintained with a force constant of 10 kcal mol^-1^ Å^-2^. In the final segment of refinement simulation, run for 250 ps at constant temperature and pressure, the same three sets of restraints listed under simulated annealing were imposed, with force constants of 5 kcal mol^-1^ Å^-2^ for backbone hydrogen bonding and for interhelical distances and 5 kcal mol^-1^ rad^-2^ for phi and psi angles (minimum at -60° for phi and -45° for psi).

The time step was 1 fs in all the simulations. All bonds connected to hydrogens were constrained by the SHAKE algorithm.^52^ Long-range electrostatic interactions were treated by the particle mesh Ewald method.^53^ The cutoff distance for nonbonded interaction was 12 Å, with force switching at 10 Å for van der Waals interactions. The Langevin thermostat with a damping constant of 1 ps^-1^ was used to maintain the temperature at 310 K and the Langevin piston^54^ was used to maintain the pressure at 1 atm.

## Acknowledgments

This work was supported by National Institutes of Health Grants AI119187, GM122698, and GM118091. All NMR experiments were carried out at the National High Magnetic Field Laboratory supported by the NSF Cooperative Agreements DMR1644779 and DMR2128556 and by the State of Florida.

## Author contributions

T.A.C., R.F. and H.-X.Z. designed the research; R.Z., H.Q., and R.P. performed the research and analyzed the data; T.A.C., R.Z., and H.-X.Z. wrote the manuscript.

