## Supplementary Figures for "Dimeric Transmembrane Structure of the SARS-CoV-2 E Protein"

*Supplemental Figures*

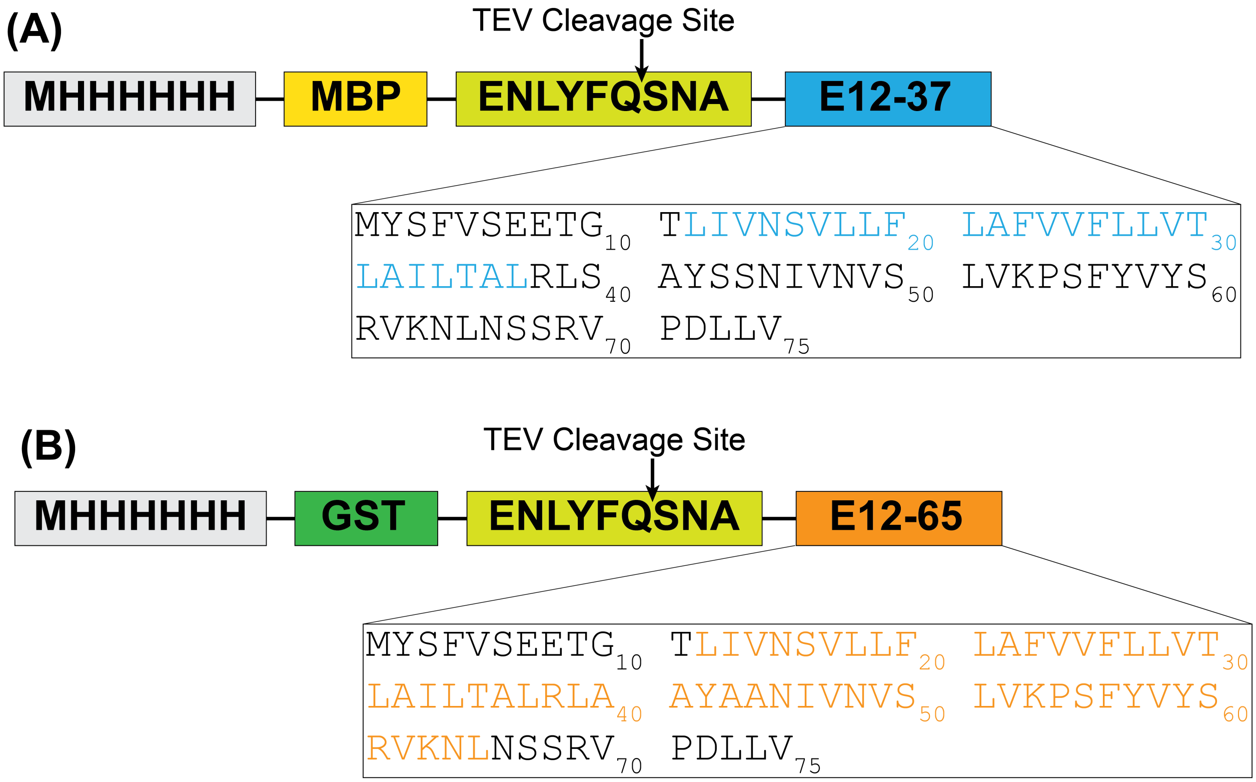


Figure S1. Schematic of SARS-CoV-2 E protein constructs utilized for bacterial expression and purification. (A) MBP fusion construct for $E_{12\text{-}37}$; $E_{1\text{-}52}$, $E_{7\text{-}43}$, and $E_{8\text{-}41}$. (B) GST fusion construct for $E_{12\text{-}65}$. The TEV cleavage site within the sequence ENLYFQSNA is indicated with an arrow.

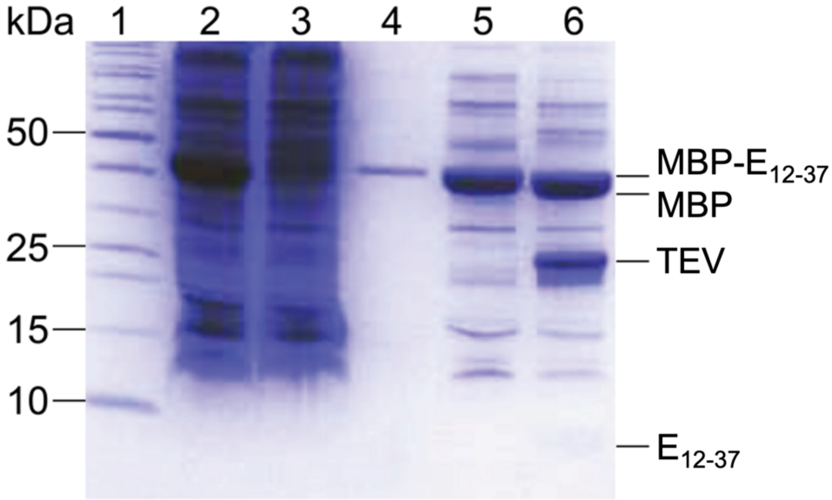


Figure S2. SDS-PAGE gel for the $E_{12\text{-}37}$ purification. Lane 1, marker; lane 2, cell lysate; lane 3, column flow-through; lane 4, column wash; lane 5, elute of MBP-$E_{12\text{-}37}$ fusion protein; lane 6, post TEV cleavage. Protein bands of interest are indicated and labeled to the right of the gel. The band for $E_{12\text{-}37}$ is actually a dimer, which could easily be mistaken as a monomer when guided only by the high molecular weight markers.

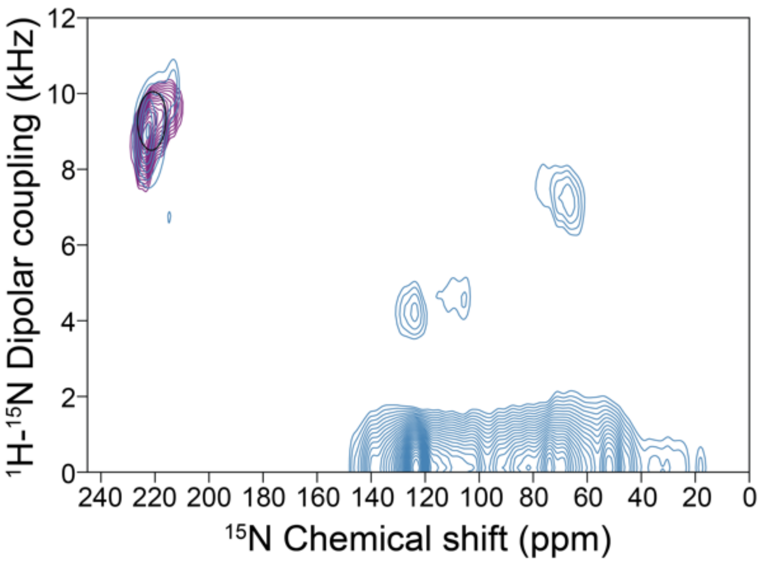


Figure S3. Overlay of 2D PISEMA spectra of $^{15}$N-Val labeled $E_{12\text{-}37}$ (purple) and $^{15}$N-Val labeled $E_{8\text{-}41}$ (teal) in aligned POPC/POPG bilayers, with a PISA wheel of 6° tilt superimposed.

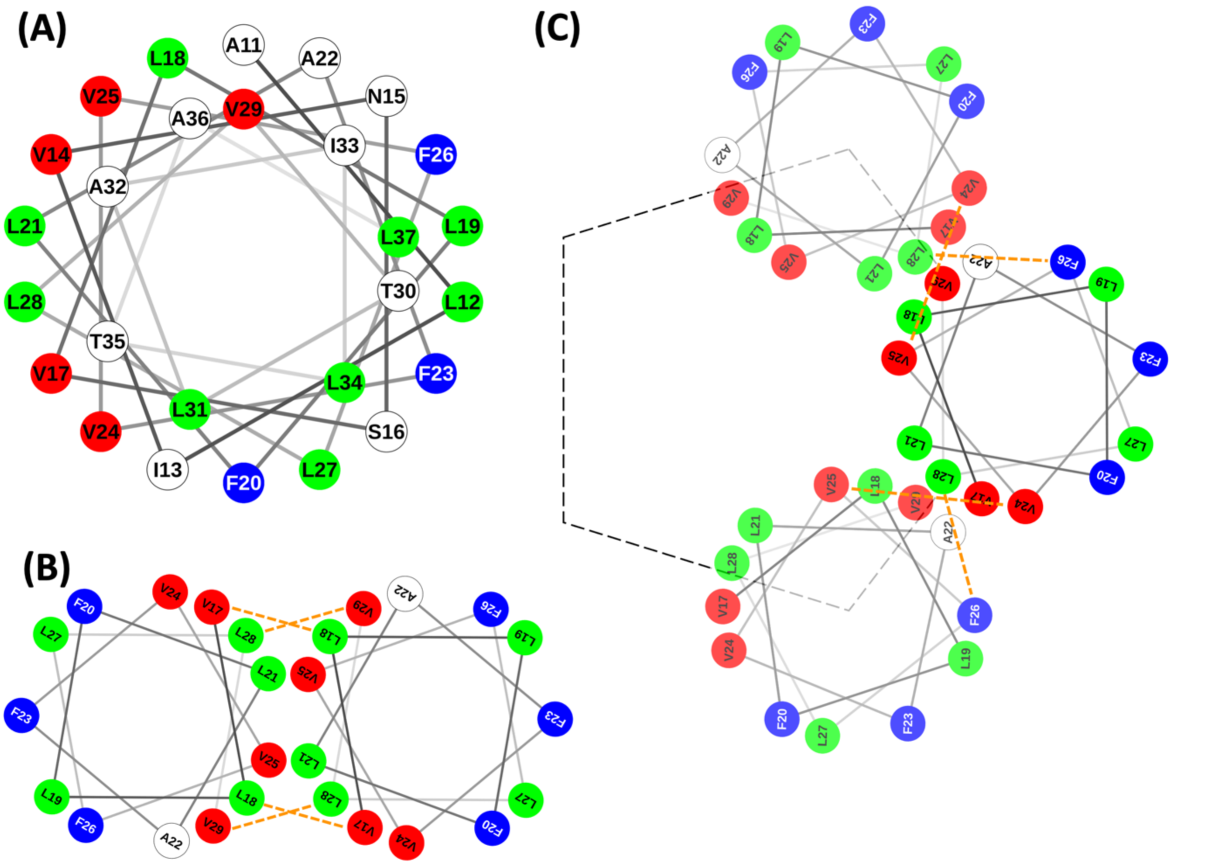


Figure S4. Possible interhelical interfaces based on the helical wheel of $E_{12\text{-}37}$. (A) The helical wheel, emphasizing the fact that five Val residues (red) locate on one face of the helix, three Phe residues (blue) locate on the opposite face of the helix, while 9 Leu residues (green) populate the entire helical surface. (B) A symmetric dimer interface that satisfies all the dipolar restraints from MAS ssNMR, including ≤ 7 Å Cα-Cα distances for all Val17-Leu18 and Leu28-Val29 pairs (indicated by dashed lines). (C) The asymmetric interhelical interface in a pentamer, minimizing the burial of Phe residues but unable to avoid Val24-Val25 (7.0 Å) and Phe26-Leu28 (7.1 Å) Cα-Cα contacts (indicated by dashed lines).

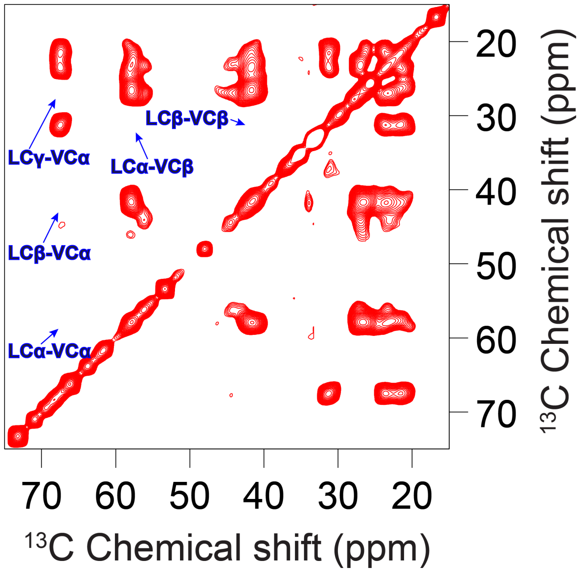


Figure S5. $^{13}C$-$^{13}C$ correlation spectrum of an equimolar mixture of $^{13}C$-Leu labeled $E_{12\text{-}37}$ and $^{13}C$-Val labeled $E_{12\text{-}37}$ in POPC/POPG liposomes at a mixing time of 100 ms. More contour levels are shown than in Figure 4 to emphasize the absence of cross peaks between labeled sites.

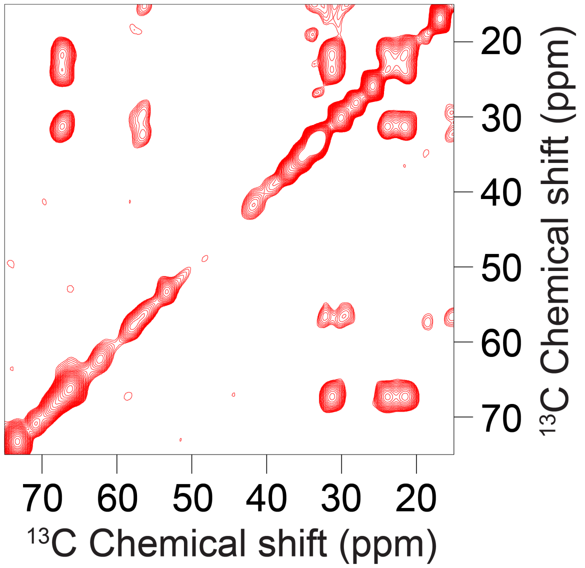


Figure S6. $^{13}C$-$^{13}C$ correlation spectrum of an equimolar mixture of two versions of the $E_{12\text{-}37}$ mutant, one with $^{13}C$-Val labeling and one with $^{13}C$-Met labeling, in POPC/POPG liposomes at a mixing time of 600 ms.

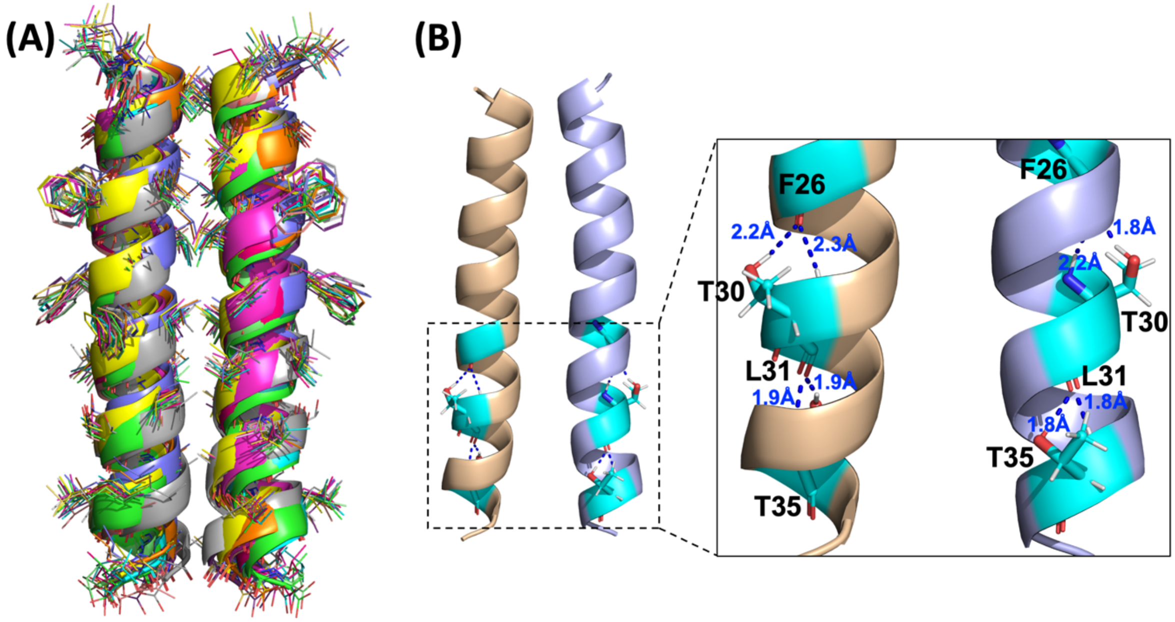


Figure S7. Refined structure of the $E_{12\text{-}37}$ dimer. (A) Superposition of 14 models from replicate refinement simulations. (B) Sidechain-backbone hydrogen bonding of Thr30 and Thr35.

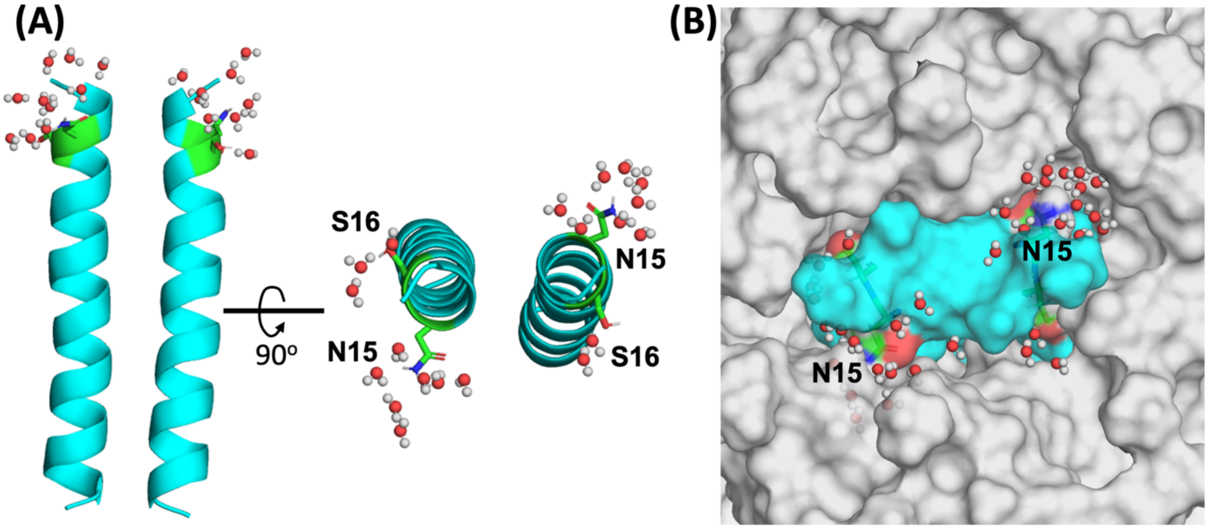


Figure S8. Exposure of Asn15 and Ser16 sidechains to water at the membrane surface. (A) Hydrogen bonding of Asn15 and Ser16 with water molecules. (B) Water-filled pockets around Asn15, potentially serving as drug-binding sites. The membrane and $E_{12\text{-}37}$ dimer are rendered as surface in gray and cyan, respectively, except that Asn15 and Ser16 are rendered with C, O, and N atoms in green, red, and blue, respectively.
